# Neural correlates of semantic number: A cross-linguistic investigation

**DOI:** 10.1101/2021.05.11.443670

**Authors:** Donald Dunagan, Shulin Zhang, Jixing Li, Shohini Bhattasali, Christophe Pallier, John Whitman, Yiming Yang, John Hale

## Abstract

One aspect of natural language comprehension is understanding how many of what or whom a speaker is referring to. While previous work has documented the neural correlates of general number comprehension and quantity comparison, we investigate semantic number from a cross-linguistic perspective with the goal of identifying cortical regions involved in distinguishing plural from singular nouns. We use three fMRI datasets in which Chinese, French, and English native speakers listen to an audiobook of a children’s story in their native language. We select these three languages because they differ in their number semantics. While Chinese lacks nominal pluralization, French and English nouns are overtly marked for number. We find a number of known semantic processing regions in common, including dorsomedial prefrontal cortex and the pars orbitalis, in which cortical activation is greater for plural than singular nouns and posit a cross-linguistic role for number in semantic comprehension.

## 1. Introduction

One aspect of natural language comprehension is understanding how many of what or whom a speaker is referring to. While much work has been done to document the neural correlates of general number comprehension and quantity comparison (e.g., Castelli et al., 2006; Dehaene et al., 2003; Kadosh & Walsh, 2009), the neural correlates of semantic number are less well understood. We use the term *semantic number* because not all of the languages which we consider morpho-syntactically mark number. More detail regarding our terminological definitions and the semantics of number is given in section 1.3. Cross-linguistically, languages with singular/plural nominal^1^ contrast tend to mark plural forms morphologically, but not mark singular forms (Corbett, 2000; Greenberg, 1963). In the semantics literature, de Swart & Farkas (2010) propose a weak singular/strong plural (unmarked singular/marked plural) account of the singular/plural contrast which respects Horn’s division of pragmatic labor (cf. Van Rooy, 2004). This is to say, in this account, plurality is morphologically and semantically marked and singularity is not. As the non-default form, we expect plural nouns to be more difficult to process than singular nouns. The first question we address is: “As measured with fMRI, do plural nouns elicit greater cortical activity than singular nouns?”

### 1.1. Number sense

One possibility which we consider is that semantic number will be subserved by the same system which subserves human *number sense*, “a short-hand for our ability to quickly understand, approximate, and manipulate numerical quantities” (Dehaene, 2001, p. 2). Natural numbers are thought to be represented as analog magnitudes along a *mental number line*. Dehaene (1992) proposes a tripartite account of number sense in the human brain. In this *triple-code* model, three portions of the parietal lobe perform different roles in number processing (Dehaene et al., 2003). The horizontal segment of the intraparietal sulcus serves as the location of the mental number line and is augmented by an angular gyrus verbal system and a posterior, superior parietal visual and attentive system.

### 1.2. Language-specific versus domain-general processing

Carreiras et al. (2010) ask if numerical processing is activated by grammatical number processing and, for stimuli with grammatical number violations, find an increase in activation in parietal regions previously implicated in number processing (Dehaene et al., 2003). Portions of the prefrontal and parietal cortices, known as the multiple-demand (MD) network (Duncan, 2010), have been found to to be responsive to a wide variety of cognitive demands such as: verbal and spatial working memory, the Stroop task, and important to this paper, an arithmetic task (Fedorenko et al., 2013). On one hand, these regions have been shown to not track linguistic input as closely as language-selective regions (Blank & Fedorenko, 2017), and Fedorenko et al. (2011) find little or no overlap between cortical regions engaged in high-level linguistic processing and MD regions which respond to general working memory, cognitive control, and importantly here, mental arithmetic. On the other hand, Carreiras et al. (2010) identify a link between number in language and general number in the brain. Our next question, then, is: “If plural nouns elicit greater cortical activity than singular nouns, do these regions of increased activation align with regions known for quantity and arithmetic processing or with regions that that are known for linguistic processing?”

### 1.3. Formal semantic motivations

Number in language is made a more interesting topic because languages can differ in their number semantics. We now review the semantics which underlie the three-way typological contrast which motivates our parallel analysis of Chinese, English, and French data. In this paper, we operate under a simplistic account of the semantics of number in which singular nominals refer to individuals and plural nominals refer to sets of individuals (but see Rullmann, 2002; de Swart & Farkas, 2010, for more detailed accounts). In the proposal by Link (1983), the entity domain to which nominals refer is a join-semilattice. Atoms are individuals and the non-atomic elements are the possible sums of multiple atoms. In these terms, singular nominals choose referents from the domain of atoms and plural nominals chose referents from the domain of sums. We also consider nominals with general number (Corbett, 2000). These are nominals which are “neutral” or “unspecified” for number. Rullmann & You (2006) describe a system for languages with general number, like Mandarin Chinese^2^, in which atoms generate a complete semilattice and nouns choose referents from the domain of atoms and sums. It is important to note that nominals with general number are not ambiguous between singular and plural readings (see Rullmann & You, 2006, for additional discussion). Their number interpretation might best be given in English as, “one or more *X*.” We distinguish semantic number from grammatical number, which is a grammatical category and which is expressed either through morphology or syntax.

A count noun (e.g., *cat*) is a noun which may be directly modified by a cardinal numerical and a mass noun (e.g., *sand*) is a noun which cannot. While (1a) is perfectly acceptable, (1b) is not acceptable on the intended reading. There is a connection between the count/mass distinction and the counting/measuring distinction. While count nouns are counted (1a), mass nouns are measured (1c). It is not the case, however, that all languages make the count/mass distinction.

(1) 
  a. two cats
  b. #two sands
  c. three buckets of sand

Chierchia (1998) proposes the Nominal Mapping Parameter, which creates a three-way typological classification for languages based upon how they express counting. Chierchia’s account is neo-Carlsonian, that is, it is based upon Carlson’s (1977) investigation of bare plurals in English. Bare nouns are nouns which occur without a determiner or a classifier. This account proposes that nouns can either be predicates at type *<e,t>*, in which they denote a set of entities, or be arguments at type *e*, in which they denote kinds. The terms *predicate* and *argument*, here, are names for the semantic types *<e,t>* (functions from individuals to truth values) and *e* (entities of argumental type), respectively. Kinds are generally understood as regularities. For the property of being a cat, there is a corresponding kind: the cat-kind. In the other direction, a kind will have a property with which it corresponds: the property of belonging to the kind.

A noun (N) may fill an argument position if it is an argument, but if it is a predicate, it must combine with a determiner to reach the argument type. Chierchia’s classification, then, is whether nouns in a language can occur as arguments, predicates, or both. From the features [+/-predicate] and [+/-argument], there are three possible language types: [+predicate, +argument], [-predicate, +argument], and [+predicate, -argument]. The type [-predicate, -argument] is not valid. English is [+predicate, +argument], Chinese is [-predicate, +argument], and French is [+predicate, -argument]. Chierchia argues that a language will have morphosyntactic properties based upon its features. The following section reviews these properties with data from Rothstein (2017, pp. 147-148).

With English being [+predicate, +argument], the nouns of English are either [+predicate] or [+argument]. Count nouns are predicates and mass nouns are arguments. Because they are predicates, singular count nouns must combine with a determiner to fill an argument position and it is predicted that bare singular count nouns are ungrammatical (2a). Plural count nouns can be shifted such that they yield a kind reading and and thus can occur as bare arguments. Mass nouns can occur bare in argument position (2b).

(2) 
  a. I saw #(a) dog.
  b. I bought wine.

Chinese allows for noun phrases (NPs) consisting of bare nouns without classifiers, number morphemes, or other functional elements. Like other classifier languages, Chinese is [-predicate, +argument]. In these languages, bare nouns can occur as arguments (3). While nouns may occur bare, they may not be directly modified by cardinal numericals. Instead of directly taking bare nouns as complements, numericals take classifier (Cl) + N sequences (4).

(3) 
  a. wǒ kànjiàn gǒu le. I see dog PART^3^ ‘I saw a dog/dogs, the dog(s).’
  b. wǒ mǎi le jiǔ. I buy PFV^4^ wine ‘I bought wine.’

(4) 
  a. sān #(zhī) gǒu three Cl_small animal_ dog ‘three dogs’
  b. liǎng #(kē) sh ù two Cl_plant_ tree ‘two trees’

In an analysis of bare noun phrases in Chinese, Yang (2001) identifies the same readings identified by Carlson (1977) for English bare plurals: kind, generic, and narrowest-scope indefinite. While French and English necessarily mark definite NPs with determiners, Chinese does not have determiners and bare NPs have definite readings that are not available in English. Since all nouns have the same properties, and no N can be directly modified by a numeral, there is no clear way to differentiate mass and count nouns grammatically. As compared to languages with mass/count distinction ([+predicate, +/-argument]), Chierchia’s (1998) view is that in [-predicate, +argument] languages, every lexical noun is mass-like. Because the plural operator does not apply to kind or mass terms, classifier languages do not have nominal pluralization. Bare nouns in these languages have a number interpretation which is general and includes the plural (3a).

French, like other Romance languages, is [+predicate, -argument] and makes the count/mass distinction. Count nouns will be marked either singular or plural and all nouns (both count and mass) must occur with a determiner (5).

(5) 
  a. J’ai vu #(un) chien. I AUX^5^ saw a dog ‘I saw a dog.’
  b. J’ai achet é #(du) vin. I AUX bought some wine ‘I bought (some) wine.’

Interestingly, French is slightly more strict with its determiner requirement than Spanish and Italian which allow for bare plurals in well-governed conditions such as object (but not subject) position. The allowed bare plurals do not have kind or generic readings, though. Because English is [+predicate, +argument], its count nouns are similar to French nouns and its mass nouns are similar to Chinese nouns.

### 1.4. Neural, cross-linguistic similarities and differences

While neural, cross-linguistic differences have been found in domains such as phonological access in a reading task (Paulesu et al., 2000), pitch contour processing (Gandour et al., 2003), and nominal and verbal representation (Li et al., 2004), similarities have been found for syntactic processing (see Obleser et al., 2011; Pallier et al., 2011, for German and French results, respectively) and comprehending linguistic content (Honey et al., 2012). Our final question, then, is: “Although they differ in their number semantics, if French, English, and Chinese display increased activation for plural nouns over singular nouns, does that activation occur in the same or different regions?”

## 2. Data and Methods

### 2.1. Participants

The English dataset includes 51 healthy, right-handed, young adults (32 female, mean age = 21.3, range = 18-37). They self-identified as native English speakers, and had no history of psychiatric, neurological or other medical illness that could compromise cognitive functions. All participants were paid and gave written informed consent prior to participation, in accordance with the guidelines of the Human Research Participant Protection Program at Cornell University.

The French dataset includes 30 healthy, right-handed adults (age range = 20-40). They self-identified as native French speakers and had no history of psychiatric, neurological, or other medical illness that could compromise cognitive functions. All participants gave written informed consent prior to participation, in accordance with the guidelines of the Regional Committee for the Protection of Persons involved in Biomedical Research.

The Chinese dataset includes 35 healthy, right-handed, young adults (15 female, mean age = 19.3, range = 18-25). They self-identified as native Chinese speakers and had no history of psychiatric, neurological, or other medical illness that could compromise cognitive functions. All participants were paid and gave written informed consent prior to participation, in accordance with the guidelines of the Ethics Committee at Jiangsu Normal University.

### 2.2. Stimuli

The French audio stimulus is an audiobook version of *Le Petit Prince* (*The Little Prince*, de Saint-Exupéry, 1946), read by Nadine Eckert-Boulet. The English audio stimulus is an English translation of *The Little Prince*, read by Karen Savage. The Chinese audio stimulus is a Chinese translation of *The Little Prince*, read by a professional female Chinese broadcaster.

The English, French, and Chinese audiobooks are 94, 98, and 99 minutes in length, respectively. The presentations were divided into nine sections, each lasting around ten minutes. Participants listened passively to the nine sections and completed four quiz questions after each section (36 questions in total). These questions were used to confirm participant comprehension of the story.

### 2.3. Data collection and preprocessing

The English and Chinese brain imaging data were acquired with a 3T MRI GE Discovery MR750 scanner with a 32-channel head coil. Anatomical scans were acquired using a T1-weighted volumetric magnetization prepared rapid gradient-echo pulse sequence. Blood-oxygen-level-dependent (BOLD) functional scans were acquired using a multi-echo planar imaging sequence with online reconstruction (TR = 2000 ms; TE’s = 12.8, 27.5, 43 ms; F = 77^*°*^; matrix size = 72 x 72; FOV = 240.0 mm x 240.0 mm; 2 x image acceleration; 33 axial slices, voxel size = 3.75 x 3.75 x 3.8 mm).

The English and Chinese fMRI data were preprocessed using AFNI version 16 (Cox, 1996). The first 4 volumes in each run were excluded from analyses to allow for T1-equilibration effects. Multi-echo independent components analysis (ME-ICA, Kundu et al., 2012) was used to denoise data for motion, physiology, and scanner artifacts. Images were then spatially normalized to the standard space of the Montreal Neurological Institute (MNI) atlas, yielding a volumetric time series resampled at 2 mm cubic voxels.

The French brain imaging data was collected with a Siemens Prisma Fit 3T scanner. T1-weighted anatomical images were acquired with a 1 mm isotropic resolution. The EPI functional images were acquired with a resolution of 3.75 x 3.75 x 3.8 mm (34 axial slices with an interleaved acquisition scheme). The 3 echo times were 10 ms, 25 ms, and 38 ms. Preprocessing was performed with ME-ICA (Kundu et al., 2012) using the default parameters and spatial normalisation was done in MNI space. The final volumetric time series consists of 3.15 mm cubic voxels.

### 2.4. Observations of interest

In order to control for discourse factors which could modulate neural activity during naturalistic language processing, we align the storybook texts and select only parallel nouns for analysis, that is, nouns which occur in all three stories and in the same context. The first step in this process is aligning sentences, which is done with the Hunalign bilingual sentence aligner (Varga et al., 2007). The alignments were checked and corrected by hand. Next, we identify the parallel nouns and filter the triplets with criteria which serve to maximize typological contrast between French, English, and Chinese nominals.

For the Chinese observations, we include only nouns which have no overt number marking, either morphological or through a number and classifier construction. This captures the [+argument] aspect of Chinese. For the French observations, we include only count nouns indexed by the definite, common determiners: *le, la, l’*, and *les*. This captures the [+predicate] aspect of French and its requirement that definite nouns be marked with a definite determiner. For the English observations, we include only count nouns, but allow them to be definite, indefinite, or type-shifted bare plurals. This capture the [+predicate, +argument] aspects of English.

While number annotation can be automated for the French nouns: *le, la*, and *l’* are singular and *les* is plural, and English count nouns are easily annotated for number based upon their overt number morphology, annotation for the Chinese nouns is more challenging because number is not overtly marked. Recall that Chinese bare nominals are not ambiguous between singular and plural readings, but it is possible that different listeners will have different judgements. Because of this, we have two native Chinese speakers annotate the Chinese nouns with singular/plural judgments. Calculating Cohen’s kappa coefficient (Cohen, 1960), a measure of inter-rater reliability, over the Chinese annotations results in a kappa = 0.96, a high degree of inter-rater reliability. We do not use any nouns in the analysis for which the two annotators disagreed in their number judgements.

The time resolution for all three of the fMRI data sets is 2.0 seconds, much slower than a natural speech rate. Because of this, we remove observations where nouns of different number would occur together within the same volume. That is, if more than one singular noun occur in the same volume or if more than one plural noun occur in the same volume, they are retained. If a singular and plural noun occur in the same volume, however, they are not kept for analysis. After this, we end up with 274 parallel observations: 245 singular and 29 plural in the Chinese text, 245 singular and 29 plural in the French text, and 244 singular and 30 plural in the English text.

### 2.5. Statistical analyses

We run separate English, French, and Chinese general linear model (GLM) analyses using Nilearn (Abraham et al., 2014; Pedregosa et al., 2011, version 0.7.1). For the English and Chinese analyses, at the first level, we include binary regressors for singular and plural nouns as well as coregressors of non-interest for spoken word rate, log lexical frequency, root mean squared amplitude of the spoken narration (RMS), and speaker pitch. Speaker pitch is not available for the French audiobook, but all of the other coregressors are used. The coregressors are added to ensure that any results found are due to the differences between singular and plural nouns and not just effects of spoken language comprehension (cf. Bullmore et al. 1999; Lund et al. 2006). The singular and plural noun regressors are marked with a 1 at the offset of the nouns of interest, word rate is marked with a 1 at the offset of every word, except for the observations of interest, log lexical frequency is marked at the end of every word, and RMS and pitch are marked every 10 ms.

Following the weak singular/strong plural (unmarked singular/marked plural) semantic account of number proposed by de Swart & Farkas (2010) (at least for French and English which make this distinction), our first-level contrast subtracts activity associated with singular nouns away from activity associated with plural nouns. At the second level, the first level contrast maps are used to perform one-sample t-tests. We apply an 8 mm full width at half maximum Gaussian smoothing kernel to counteract inter-subject anatomical variation. The by language, group-level results reported in the following section underwent family-wise-error (FWE) voxel correction for multiple comparisons and are reported in terms of z-score. We only retain clusters greater than 100 mm^3^. Additionally, because we want to identify any common regions of increased activation between the three languages, we report the overlap of the separate results. The MNI2TAL tool from the Yale BioImage Suite^6^ (Lacadie et al., 2008a,b, version 1.4) was referenced for brain region and Brodmann area labels.

## 3. Results

### 3.1. Chinese results

For the Chinese participants, we find an increase in activation for plural nouns over singular nouns in the left pars triangularis (BA 45) and left pars orbitalis (BA 47), extending into left dorsolateral prefrontal corex (BA 46), in left dorsomedial prefrontal cortex (BAs 8, 9), the left fusiform (BA 37), angular (BA 39), and middle temporal (BA 21) gyri, and the right cerebellum. These results can be seen in Fig. 1a and more detail can be found in Table 1.

**Table 1:** Significant PLURAL > SINGULAR clusters for Chinese after FWE voxel correction for multiple comparisons with p < 0.05, cluster size > 100 mm^3^.

| Region | Cluster size<br>(mm<sup>3</sup>) | MNI coordinates |  |  | Peak stat (z) |
| --- | --- | --- | --- | --- | --- |
|  |  | x | y | z |  |
| L Pars Orbitalis (BA 47) | 4936 | -40.0 | 38.0 | -16.0 | 6.85 |
| L Dorsolateral Prefrontal Cortex (BA 46) |  | -46.0 | 42.0 | 2.0 | 5.62 |
| L Dorsomedial Prefrontal Cortex (BA 9) | 6952 | -12.0 | 44.0 | 38.0 | 6.70 |
| L Dorsomedial Prefrontal Cortex (BA 8) |  | -10.0 | 30.0 | 48.0 | 6.45 |
|  |  | -8.0 | 20.0 | 56.0 | 5.64 |
| Left Fusiform Gyrus (BA 37) | 768 | -60.0 | -42.0 | -12.0 | 5.54 |
| Left Angular Gyrus (BA 39) | 304 | -36.0 | -64.0 | 26.0 | 5.54 |
| Left Middle Temporal Gyrus (BA 21) | 192 | -56.0 | -6.0 | -22.0 | 5.44 |
| L Pars Triangularis (BA 45) | 208 | -46.0 | 28.0 | 4.0 | 5.39 |
| R Cerebellum | 136 | 36.0 | -76.0 | -36.0 | 5.39 |
| L Angular Gyrus (BA 39) | 176 | -34.0 | -78.0 | 44.0 | 5.33 |
| L Dorsal Prefrontal Cortex (BA 8) | 104 | -26.0 | 18.0 | 56.0 | 5.21 |

**Figure 1:**
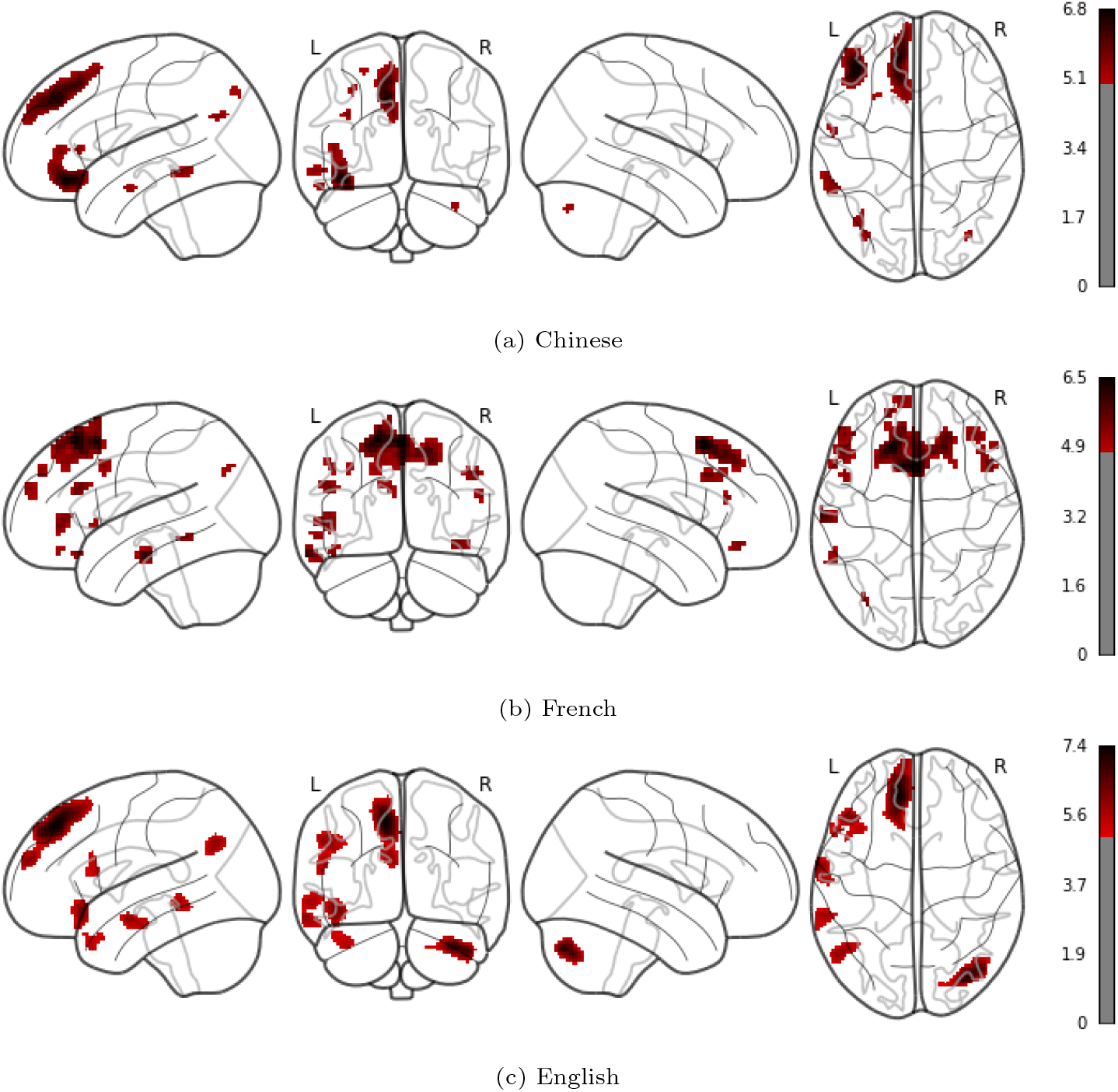
Significant clusters for PLURAL > SINGULAR contrast (z-valued) after FWE voxel correction for multiple comparisons with p < 0.05, cluster size > 100 mm^3^ for Chinese (a), French (b), and English (c).

### 3.2. French results

For the French participants, we find an increase in activation for plural nouns over singular nouns in the left and right pars opercularis (BA 44), left pars triangularis (BA 45), left and right pars orbitalis (BA 47), left (BAs 8, 9, 10) and right (BA 8) dorsomedial prefrontal cortex, left (BAs 8, 46) and right (BA 9) dorsolateral prefrontal cortex, and the left middle temporal (BA 21), fusiform (BA 37), and angular (BA 39) gyri. These results can be seen in Fig. 1b and more detail can be found in Table 2.

**Table 2:** Significant PLURAL > SINGULAR clusters for French after FWE voxel correction for multiple comparisons with p < 0.05, cluster size > 100 mm^3^.

| Region | Cluster size<br>(mm<sup>3</sup>) | MNI coordinates |  |  | Peak stat (z) |
| --- | --- | --- | --- | --- | --- |
|  |  | x | y | z |  |
| Dorsomedial Prefrontal Cortex (BA 8) | 14917 | -14.0 | 30.0 | 60.0 | 6.48 |
|  |  | 2.0 | 15.0 | 56.0 | 6.45 |
|  |  | -8.0 | 18.0 | 56.0 | 6.34 |
|  |  | 17.0 | 30.0 | 50.0 | 5.91 |
| L Middle Temporal Gyrus (BA 21) | 1040 | -65.0 | -17.0 | -19.0 | 6.07 |
| R Pars Opercularis (BA 44) | 914 | 52.0 | 18.0 | 34.0 | 5.66 |
| R Dorsolateral Prefrontal Cortex (BA 9) |  | 46.0 | 27.0 | 34.0 | 5.25 |
| L Pars Opercularis (BA 44) | 1040 | -49.0 | 27.0 | 25.0 | 5.62 |
| L Dorsolateral Prefrontal Cortex (BA 46) | 946 | -49.0 | 40.0 | 3.0 | 5.43 |
| R Pars Orbitalis (BA 47) | 630 | 39.0 | 40.0 | -13.0 | 5.38 |
| L Dorsolateral Prefrontal Cortex (BA 8) | 473 | -49.0 | 11.0 | 41.0 | 5.29 |
| R Dorsolateral Prefrontal Cortex (BA 9) | 220 | 55.0 | 30.0 | 18.0 | 5.27 |
| L Dorsomedial Prefrontal Cortex (BA 10) | 693 | -8.0 | 62.0 | 25.0 | 5.26 |
| L Pars Orbitalis (BA 47) | 157 | -46.0 | 30.0 | -19.0 | 5.23 |
| L Dorsomedial Prefrontal Cortex (BA 9) | 378 | -17.0 | 56.0 | 37.0 | 5.21 |
| L Fusiform Gyrus (BA 37) | 378 | -58.0 | -45.0 | -10.0 | 5.13 |
| L Pars Triangularis (BA 45) | 189 | -55.0 | 18.0 | -0.0 | 5.04 |
| L Angular Gyrus (BA 39) | 220 | -33.0 | -74.0 | 41.0 | 5.03 |
| L Pars Orbitalis (BA 47) | 157 | -49.0 | 43.0 | -19.0 | 5.01 |

### 3.3. English results

For the English participants, we find an increase in activation for plural nouns over singular nouns in the left pars opercularis (BA 44), left pars triangularis (BA 45), left pars orbitalis (BA 47), left dorsomedial prefrontal cortex (BAs 8, 10), the left temporal pole (BA 38), the left middle temporal (BA 21) and angular gyri (BA 39), and the the right cerebellum. These results can be seen in Fig. 1c and more detail can be found in Table 3.

**Table 3:**
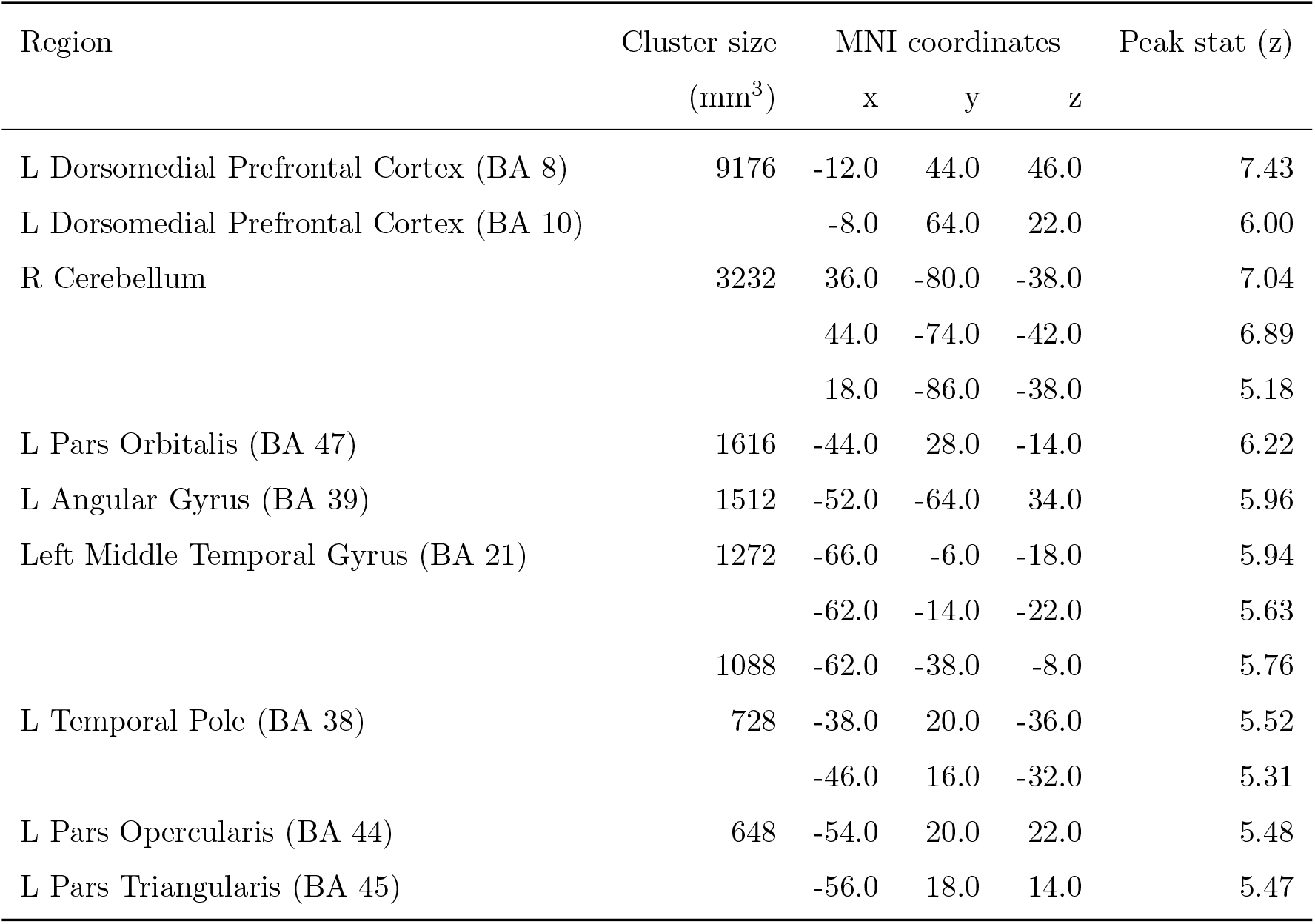
Significant PLURAL > SINGULAR clusters for English after FWE voxel correction for multiple comparisons with p < 0.05, cluster size > 100 mm^3^.

### 3.4. Cross-linguistic overlap

Overlaying the significant clusters from the Chinese, French, and English main results, we find voxel-wise overlap between all three languages in the left pars orbitalis (BA 47) and left dorsomedial prefrontal cortex (BAs 8, 10), as indicated in black in Fig. 2. We find voxel-wise overlap between two languages in the left pars orbitalis (BA 47), left dorsomedial prefrontal cortex (BA 8), left dorsolateral prefrontal cortex (BA 46), the left middle temporal (BA 21) and fusiform (BA 37) gyri, and the right cerebellum, as indicated in red in Fig. 2. More detail can be found in Table 4. Additionally, while we do not observe voxel-level overlap, all three languages show an increase in activation in the left pars triangularis (BA 45), and the left angular gyrus (BA 39).

**Table 4:** Clusters resulting from overlap of Chinese, French, and English PLURAL > SINGULAR main results. Only clusters where 2 or more languages overlap are presented.

| Region | Cluster size<br>(mm<sup>3</sup>) | MNI coordinates |  |  | Overlapping<br>languages |
| --- | --- | --- | --- | --- | --- |
|  |  | x | y | z |  |
| L Pars Orbitalis (BA 47) | 504 | -46.0 | 27.0 | -19.0 | 3.0 |
| L Dorsomedial Prefrontal Cortex (BA 10) | 504 | -8.0 | 62.0 | 25.0 | 3.0 |
| L Dorsomedial Prefrontal Cortex (BA 8) | 4194 | -14.0 | 30.0 | 53.0 | 3.0 |
|  |  | -11.0 | 40.0 | 47.0 | 3.0 |
|  |  | -8.0 | 24.0 | 56.0 | 3.0 |
|  |  | -21.0 | 30.0 | 53.0 | 2.0 |
| L Middle Temporal Gyrus (BA 21) | 189 | -62.0 | -14.0 | -23.0 | 2.0 |
| L Fusiform Gyrus (BA 37) | 220 | -58.0 | -42.0 | -10.0 | 2.0 |
| L Dorsolateral Prefrontal Cortex (BA 46) | 220 | -46.0 | 43.0 | 3.0 | 2.0 |
| L Pars Orbitalis (BA 47) | 126 | -46.0 | 40.0 | -16.0 | 2.0 |
| L Dorsomedial Prefrontal Cortex (BA 8) | 31 | -5.0 | 27.0 | 50.0 | 2.0 |
| R Cerebellum | 63 | 36.0 | -74.0 | -35.0 | 2.0 |

**Figure 2:**
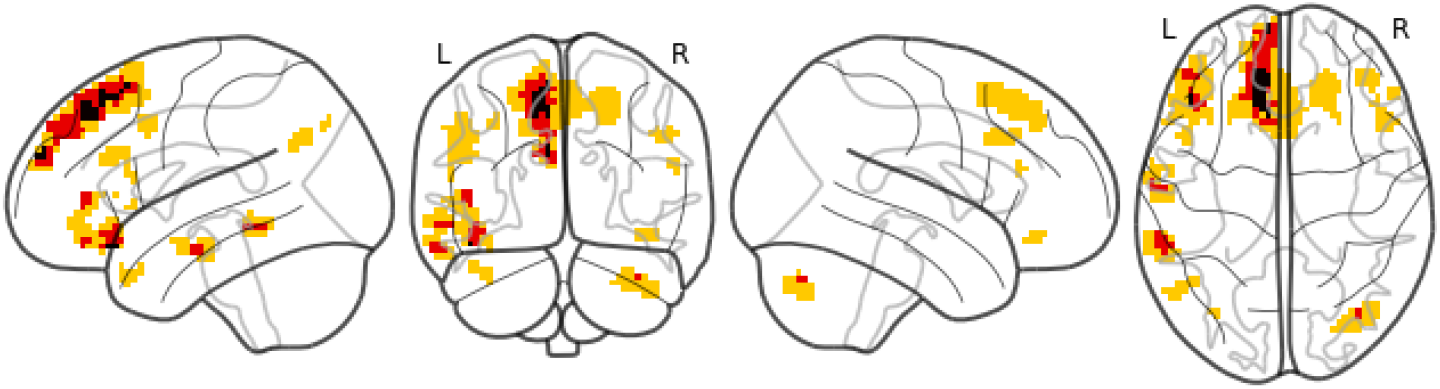
Overlap of Chinese, French, and English main results. Yellow indicates significant PLURAL > SINGULAR clusters for 1 language, red for 2 languages, and black for all 3 languages.

## 4. Discussion

In contrasting neural activation between plural and singular nouns, we observe several common regions of increased activation between the three languages: the left pars orbitalis (POrb), left pars triangularis (PTri), left dorsomedial (DMPFC) and dorsolateral (DLPFC) prefrontal cortex, the left middle temporal (MTG), fusiform, and angular gyri (AG), and the right cerebellum. We observe voxel-level overlap between all three languages in the left POrb and left DMPFC. Our findings do not align with what would be expected if semantic number were subserved by the same brain network as triple-code number processing and quantity comparison (Dehaene et al., 2003). Instead, the brain regions which we identify are more similar to those identified for semantic processing (e.g., Binder et al., 2009; Huth et al., 2016). More specifically, they align with regions previously implicated in fMRI studies of multi-word semantic comprehension (e.g., Graessner et al., 2021; Graves et al., 2010). Our interpretation of the results is that it is more difficult to process plural nouns than it is to process singular nouns.

### 4.1. Quantity comparison

The triple-code model (Dehaene, 1992) proposes a tripartite account for making sense of numbers and quantities with three portions of the parietal lobe facilitating this in different manners (Dehaene et al., 2003). The horizontal segment of the intraparietal sulcus (HIPS) serves as an internal number line which keeps track of size and distance between numbers and which is responsible for number representation, the left AG aids in processing heard numbers without processing quantities directly, and the posterior, superior parietal lobe (PSPL) orients attention both in space and on the internal number line. While the HIPS would be a plausible candidate for our PLURAL > SINGULAR contrast, a significant difference in activation is not observed there. Even though Carreiras et al. (2010) observe an increase in activation for grammatical number disagreement in the right HIPS and right PSPL, they believe it unlikely that the parietal regions are specifically involved when processing language and it more likely that the activation is from quantity computation engaged by the grammatical judgement task. Indeed, they perform SINGULAR > PLURAL and PLURAL > SINGULAR contrasts for the identified parietal regions, but find no effect. While we do identify the left AG, we, importantly, do not identify the HIPS in our contrast of plural and singular nouns. If quantity comparison specific processing were happening, we would expect to identify the HIPS.

### 4.2. Semantic processing

Binder et al. (2009) and Huth et al. (2016) demonstrate that the semantic network is both distributed and diverse. The regions which we find for our semantic number contrast are not out of place when compared to their results. However, their focus is on conceptual knowledge storage. In the case of Binder et al. (2009), the reviewed studies typically only identify one region or one of the region groups, depending on the semantic contrast. In the case of Huth et al. (2016), they find that while a concept like *self* may be selected for by portions of more than one region (e.g., PFC, AG, and MTG), those portions are small and interspersed among portions which select for other concepts in that region. In comparison, we identify a multitude of regions with our single contrast. Additionally, while our stimuli are embedded in naturalistic sentences, our analysis is not a data-driven one like that of Huth et al. (2016). The divergence of our results from theirs shows that linguistic plurality is not a concept stored in one brain region.

Another facet of semantics is multi-word comprehension. For explicit, two-word semantic composition Graessner et al. (2021) find an increase in activation in the left IFG, left DMPFC, bilateral AG, left pMTG, left ATL, right fusiform gyrus, and the right cerebellum, among other regions and Graves et al. (2010) find an increase in activation in the right AG, bilateral DMPFC, and bilateral posterior cingulate and precuneus. For semantic comprehension of sentence-length stimuli, Pallier et al. (2011) identify the left ATL, left anterior superior temporal sulcus (STS), and left temporo-parietal junction and Humphries et al. (2006) identify bilateral STS, MTG, inferior temporal gyrus, and AG (the frontal lobe was not included in their analysis). Our stimuli are in-between those of these two modalities: our observations of interest are not sentences, but neither are they two-word noun phrases. Our results are most similar to those of Graessner et al. (2021), though.

Our interpretation of this conflict is that we are tapping into a semantic representation intermediate between two-word noun phrases and full-fledged sentences and that there is an effect for nominal semantic number. That is, it is more difficult to integrate plural nouns into the current, working semantic representation than singular nouns. It is understandable that whether there are *one* or *many* of someone or something would play a role in constructing meaning during language comprehension and that being morphologically (Corbett, 2000; Greenberg, 1963) and semantically (de Swart & Farkas, 2010) marked, plural nominals would elicit greter activation than singular nominals.

It is interesting that we only identify the left ATL in the English results, but not the Chinese or French results, given its importance in the literature on semantic composition (e.g., Bemis & Pylkkänen, 2011, 2013; Li et al., 2020). The anterior temporal lobes can be susceptible to signal loss during fMRI imaging (Devlin et al., 2000), however, and we perform full-brain analyses, which reduces statistical power.

### 4.3. Similarities and differences between the results

The similarities that we see between the Chinese, English, and French results are not unprecedented. In bilinguals, previous research has found overlap between the L1 and L2 regions which subserve lexicosemantic comparison (Crinion et al., 2006; Klein et al., 2006). Honey et al. (2012) expand this to narrative level stimuli, analyzing neural activity for monolingual English speakers and bilingual, Russian native, English L2 speakers in two conditions: listening to a Russian story and listening to an English translation of that story. When the participants listen to the story in their native language, they find a number of areas in common which reliably respond to the content of the narrative: the STS, the AG, the supramarginal gyrus, the IFG, the precuneus, the middle frontal gyrus, and orbitofrontal cortex. These results, like ours, show that neural response patterns can be shared across groups despite differences in linguistic form. Importantly, we find that plurality conveyed through discourse cues (Chinese) elicits a similar response to overtly marked plurality (French and English).

With regard to the the differences between the Chinese, English, and French results, we observe some differences in the regions which show an increase in activation. Only in the English results do we identify the left ATL and the French results are much more bilateral than the English and Chinese results. Curiously, though, the French results do not implicate the right cerebellum. Some possible explanations for the observed differences include differences in salience and location of number marking. While the semantic number of our non-number marked Chinese observations is conveyed to the listener through discourse cues, for the French observations, number is overtly marked on the determiner which occurs before the noun and for the English observations, number is morphologically marked on the noun itself. Another potential factor is the differences between the three datasets. They were collected by different researchers in different facilities and the Chinese and English datasets have a higher resolution than the French dataset, which leads to a more aggressive FWE correction.

## 5. Conclusion

In this project, we investigate the neural correlates of semantic number from a cross-linguistic perspective. We find that plural nouns elicit greater cortical activity than singular nouns. This is consistent with the account of de Swart & Farkas (2010) in which plural is semantically and morphologically marked and singular is not. While Chinese does not overtly mark bare nominals for number, French and English count nouns are overtly marked for number. Despite the differences in their semantics and the resulting morphosyntactic properties, there is much overlap between the regions of the three languages in which we observe an increase in activation for plural over singular nouns. We observe voxel-level overlap in the left pars orbitalis and left dorsomedial prefrontal cortex. We discuss our findings with respect to previous cognitive neuroscience and neurolinguistic research and argue that semantic comprehension (e.g., Graessner et al., 2021; Graves et al., 2010) and not quantity comparison specific (Dehaene et al., 2003) processing is occurring when listeners process semantic number.

The neurobiology of language research domain (e.g., Bornkessel-Schlesewsky & Schlesewsky, 2016; Kemmerer, 2016; Poeppel et al., 2012) is interested in explaining how language is implemented in the human brain and one aspect that must be accounted for in any comprehensive model is the similarities and differences between languages described by research in linguistic typology. Our research advances this goal by investigating Chierchia’s (1998) typological counting distinction in which nouns can be predicates, arguments, or both, using French, Chinese, and English, as representative languages.

## Footnotes

1 Nominal is used as a cover term for noun phrases and determiner phrases.

2 From here forward, we will simply say Chinese.

3 Particle

4 Perfective

5 Auxiliary

6 https://bioimagesuiteweb.github.io/webapp/mni2tal.html

## References

Abraham, A., Pedregosa, F., Eickenberg, M., Gervais, P., Mueller, A., Kossaifi, J., Gramfort, A., Thirion, B., & Varoquaux, G. (2014). Machine learning for neuroimaging with scikit-learn. Frontiers in Neuroinformatics, 8, 14.

Bemis, D. K., & Pylkkänen, L. (2011). Simple composition: A magnetoencephalography investigation into the comprehension of minimal linguistic phrases. Journal of Neuroscience, 31, 2801–2814.

Bemis, D. K., & Pylkkänen, L. (2013). Basic linguistic composition recruits the left anterior temporal lobe and left angular gyrus during both listening and reading. Cerebral Cortex, 23, 1859–1873.

Binder, J. R., Desai, R. H., Graves, W. W., & Conant, L. L. (2009). Where is the semantic system? a critical review and meta-analysis of 120 functional neuroimaging studies. Cerebral Cortex, 19, 2767–2796.

Blank, I. A., & Fedorenko, E. (2017). Domain-general brain regions do not track linguistic input as closely as language-selective regions. Journal of Neuroscience, 37, 9999–10011.

Bornkessel-Schlesewsky, I., & Schlesewsky, M. (2016). The importance of linguistic typology for the neurobiology of language. Linguistic Typology, 20, 615–621.

Bullmore, E. T., Brammer, M. J., Rabe-Hesketh, S., Curtis, V. A., Morris, R. G., Williams, S. C., Sharma, T., & McGuire, P. K. (1999). Methods for diagnosis and treatment of stimulus-correlated motion in generic brain activation studies using fMRI. Human Brain Mapping, 7, 38–48.

Carlson, G. N. (1977). Reference to kinds in English.. Ph.D. thesis University of Massachusetts at Amherst.

Carreiras, M., Carr, L., Barber, H. A., & Hernandez, A. (2010). Where syntax meets math: Right intraparietal sulcus activation in response to grammatical number agreement violations. NeuroImage, 49, 1741–1749.

Castelli, F., Glaser, D. E., & Butterworth, B. (2006). Discrete and analogue quantity processing in the parietal lobe: A functional mri study. Proceedings of the National Academy of Sciences, 103, 4693–4698.

Chierchia, G. (1998). Reference to kinds across language. Natural Language Semantics, 6, 339–405.

Cohen, J. (1960). A coefficient of agreement for nominal scales. Educational and Psychological Measurement, 20, 37–46.

Corbett, G. G. (2000). Number. Cambridge: Cambridge University Press.

Cox, R. W. (1996). Afni: software for analysis and visualization of functional magnetic resonance neuroimages. Computers and Biomedical Research, 29, 162–173.

Crinion, J., Turner, R., Grogan, A., Hanakawa, T., Noppeney, U., Devlin, J. T., Aso, T., Urayama, S., Fukuyama, H., Stockton, K. et al. (2006). Language control in the bilingual brain. Science, 312, 1537–1540.

Dehaene, S. (1992). Varieties of numerical abilities. Cognition, 44, 1–42.

Dehaene, S. (2001). Précis of the number sense. Mind & Language, 16, 16–36.

Dehaene, S., Piazza, M., Pinel, P., & Cohen, L. (2003). Three parietal circuits for number processing. Cognitive Neuropsychology, 20, 487–506.

Devlin, J. T., Russell, R. P., Davis, M. H., Price, C. J., Wilson, J., Moss, H. E., Matthews, P. M., & Tyler, L. K. (2000). Susceptibility-induced loss of signal: comparing pet and fmri on a semantic task. Neuroimage, 11, 589–600.

Duncan, J. (2010). The multiple-demand (md) system of the primate brain: mental programs for intelligent behaviour. Trends in Cognitive Sciences, 14, 172–179.

Fedorenko, E., Behr, M. K., & Kanwisher, N. (2011). Functional specificity for high-level linguistic processing in the human brain. Proceedings of the National Academy of Sciences, 108, 16428–16433.

Fedorenko, E., Duncan, J., & Kanwisher, N. (2013). Broad domain generality in focal regions of frontal and parietal cortex. Proceedings of the National Academy of Sciences, 110, 16616–16621.

Gandour, J., Dzemidzic, M., Wong, D., Lowe, M., Tong, Y., Hsieh, L., Sattham-nuwong, N., & Lurito, J. (2003). Temporal integration of speech prosody is shaped by language experience: An fmri study. Brain and Language, 84, 318–336.

Graessner, A., Zaccarella, E., & Hartwigsen, G. (2021). Differential contributions of left-hemispheric language regions to basic semantic composition. Brain Structure and Function, 226, 501–518.

Graves, W. W., Binder, J. R., Desai, R. H., Conant, L. L., & Seidenberg, M. S. (2010). Neural correlates of implicit and explicit combinatorial semantic processing. Neuroimage, 53, 638–646.

Greenberg, J. H. (1963). Universals of language.. Cambridge, MA: MIT Press.

Honey, C. J., Thompson, C. R., Lerner, Y., & Hasson, U. (2012). Not lost in translation: neural responses shared across languages. Journal of Neuroscience, 32, 15277–15283.

Humphries, C., Binder, J. R., Medler, D. A., & Liebenthal, E. (2006). Syntactic and semantic modulation of neural activity during auditory sentence comprehension. Journal of Cognitive Neuroscience, 18, 665–679.

Huth, A. G., De Heer, W. A., Griffiths, T. L., Theunissen, F. E., & Gallant, J. L. (2016). Natural speech reveals the semantic maps that tile human cerebral cortex. Nature, 532, 453–458.

Kadosh, R. C., & Walsh, V. (2009). Numerical representation in the parietal lobes: Abstract or not abstract? Behavioral and Brain Sciences, 32, 313–328.

Kemmerer, D. (2016). Do language-specific word meanings shape sensory and motor brain systems? the relevance of semantic typology to cognitive neuroscience. Linguistic Typology, 20, 623–634.

Klein, D., Zatorre, R. J., Chen, J.-K., Milner, B., Crane, J., Belin, P., & Bouffard, M. (2006). Bilingual brain organization: A functional magnetic resonance adaptation study. Neuroimage, 31, 366–375.

Kundu, P., Inati, S. J., Evans, J. W., Luh, W.-M., & Bandettini, P. A. (2012). Differentiating bold and non-bold signals in fmri time series using multi-echo epi. Neuroimage, 60, 1759–1770.

Lacadie, C., Fulbright, R., Arora, J., Constable, R., & Papademetris, X. (2008a). Brodmann areas defined in mni space using a new tracing tool in bioimage suite. In Proceedings of the 14th annual meeting of the organization for human brain mapping. volume 771.

Lacadie, C. M., Fulbright, R. K., Rajeevan, N., Constable, R. T., & Papademetris, X. (2008b). More accurate talairach coordinates for neuroimaging using non-linear registration. Neuroimage, 42, 717–725.

Li, J. et al. (2020). Disentangling semantic composition and semantic association in the left temporal lobe. bioRxiv,.

Li, P., Jin, Z., & Tan, L. H. (2004). Neural representations of nouns and verbs in chinese: an fmri study. Neuroimage, 21, 1533–1541.

Link, G. (1983). The logical analysis of plurals and mass terms: A lattice-theoretical approach. Formal Semantics: The Essential Readings, 127, 147.

Lund, T. E., Madsen, K. H., Sidaros, K., Luo, W.-L., & Nichols, T. E. (2006). Non-white noise in fMRI: does modelling have an impact? Neuroimage, 29, 54–66.

Obleser, J., Meyer, L., & Friederici, A. D. (2011). Dynamic assignment of neural resources in auditory comprehension of complex sentences. Neuroimage, 56, 2310–2320.

Pallier, C., Devauchelle, A.-D., & Dehaene, S. (2011). Cortical representation of the constituent structure of sentences. Proceedings of the National Academy of Sciences, 108, 2522–2527.

Paulesu, E., McCrory, E., Fazio, F., Menoncello, L., Brunswick, N., Cappa, S. F., Cotelli, M., Cossu, G., Corte, F., Lorusso, M. et al. (2000). A cultural effect on brain function. Nature Neuroscience, 3, 91–96.

Pedregosa, F., Varoquaux, G., Gramfort, A., Michel, V., Thirion, B., Grisel, O., Blondel, M., Prettenhofer, P., Weiss, R., Dubourg, V. et al. (2011). Scikit-learn: Machine learning in python. the Journal of Machine Learning Research, 12, 2825–2830.

Poeppel, D., Emmorey, K., Hickok, G., & Pylkkänen, L. (2012). Towards a new neurobiology of language. Journal of Neuroscience, 32, 14125–14131.

Rothstein, S. (2017). Semantics for counting and measuring. Cambridge: Cambridge University Press.

Rullmann, H. (2002). Bound-variable pronouns and the semantics of number. In Proceedings of the western conference on linguistics: Wecol (pp. 243–254). volume 14.

Rullmann, H., & You, A. (2006). General number and the semantics and pragmatics of indefinite bare nouns in mandarin chinese. Where Semantics meets Pragmatics, (pp. 175–196).

de Saint-Exupéry, A. (1946). Le petit prince [The little prince]. Paris: Gallimard.

de Swart, H., & Farkas, D. (2010). The semantics and pragmatics of plurals. Semantics and Pragmatics 3, 6–1.

Van Rooy, R. (2004). Signalling games select horn strategies. Linguistics and Philosophy, 27, 493–527.

Varga, D., Halácsy, P., Kornai, A., Nagy, V., Németh, L., & Trón, V. (2007). Parallel corpora for medium density languages. Amsterdam Studies In The Theory And History Of Linguistic Science Series 4, 292, 247.

Yang, R. (2001). Common nouns, classifiers, and quantification in Chinese. Ph.D. thesis Rutgers University.

